# Ketamine rescues anhedonia by cell-type and input specific adaptations in the Nucleus Accumbens

**DOI:** 10.1101/2023.06.08.544088

**Authors:** Federica Lucantonio, Shuwen Li, Jaden Lu, Jacob Roeglin, Leonardo Bontempi, Brenda C. Shields, Carlos A. Zarate, Michael R. Tadross, Marco Pignatelli

## Abstract

Ketamine’s role in providing a rapid and sustained antidepressant response, particularly for patients unresponsive to conventional treatments, is increasingly recognized. A core symptom of depression, anhedonia, or the loss of enjoyment or interest in previously pleasurable activities, is known to be significantly alleviated by ketamine. While several hypotheses have been proposed regarding the mechanisms by which ketamine alleviates anhedonia, the specific circuits and synaptic changes responsible for its sustained therapeutic effects are not yet understood. Here, we show that the nucleus accumbens (NAc), a major hub of the reward circuitry, is essential for ketamine’s effect in rescuing anhedonia in mice subjected to chronic stress, a critical risk factor in the genesis of depression in humans. Specifically, a single exposure to ketamine rescues stress-induced decreased strength of excitatory synapses on NAc D1 dopamine receptor-expressing medium spiny neurons (D1-MSNs). By using a novel cell-specific pharmacology method, we demonstrate that this cell-type specific neuroadaptation is necessary for the sustained therapeutic effects of ketamine. To test for causal sufficiency, we artificially mimicked ketamine-induced increase in excitatory strength on D1-MSNs and found that this recapitulates the behavioral amelioration induced by ketamine. Finally, to determine the presynaptic origin of the relevant glutamatergic inputs for ketamine-elicited synaptic and behavioral effects, we used a combination of opto- and chemogenetics. We found that ketamine rescues stress-induced reduction in excitatory strength at medial prefrontal cortex and ventral hippocampus inputs to NAc D1-MSNs. Chemogenetically preventing ketamine-evoked plasticity at those unique inputs to the NAc reveals a ketamine-operated input-specific control of hedonic behavior. These results establish that ketamine rescues stress-induced anhedonia via cell-type-specific adaptations as well as information integration in the NAc via discrete excitatory synapses.

---

Depression is a significant public health concern that must be addressed urgently in order to reduce the burden of disease and disability^1–3^. However, current pharmacotherapies for depression are associated with a high non-response rate and require prolonged administration for clinical improvement, often lasting weeks or even months^4, 5^. In contrast, a single sub-anesthetic dose of ketamine, a noncompetitive N-methyl-D-aspartate (NMDA) glutamate receptor antagonist known to lead to an amplification of α-amino-3-hydroxy-5-methyl-4-isoxazole propionic acid receptor (AMPAR)-mediated excitatory synaptic transmission^6–8^, has been shown to induce a rapid and sustained antidepressant effect within a matter of hours in 70% of treatment-resistant patients^7, 9–13^. Importantly, ketamine reduces anhedonia^14–17^, which is one of the two core symptoms in the diagnosis of major depression and is defined as a diminished pleasure or interest in previously rewarding activities^18–20^. Notably, conventional medications for depression are often inadequate in alleviating anhedonia^21^.

Despite these promising results, the neurobiological mechanisms of ketamine’s anti-anhedonic action remain unclear. Because the antidepressant impact of ketamine persists long after its metabolic breakdown^22, 23^, its mechanism of action may result from long-lasting adaptations in neural circuits implicated in motivated behavior. However, the identity of these circuits and the sustained adaptations they may undergo upon treatment with ketamine have not been systematically investigated, and our ability to develop more targeted, effective, and safer therapies is therefore limited. To that end, here we provide a comprehensive framework for understanding the circuit and synaptic bases of ketamine’s sustained alleviation of anhedonia using mice as a model system.

## Ketamine rescues stress-induced anhedonia

Mice assign hedonic value to a variety of stimuli that also elicit hedonic value in humans, such as palatable foods (e.g. sweets, like sucrose), and social interactions^18^. Chronic stress, a crucial risk factor in the genesis of depression in humans, induces hedonic deficits in rodents^24–26^. Specifically, chronic stress in mice causes deficits in the two main components of anhedonia: consummatory (subjective pleasure) and motivational (anticipation of and drive towards rewarding stimuli) behavior^18^. Here, we exposed mice to chronic corticosterone (CORT), the main murine stress hormone that mediates the effects of chronic stress on behavior^27–31^, and tested its effect on both consummatory and motivational components of anhedonia by using a battery of behavioral tests, including sucrose preference test, three-chamber sociability test, and sucrose self-administration procedure (Fig. 1a,e; Extended Data Fig. 1). Mice subjected to chronic CORT administration (21 days) displayed a reduction in sucrose preference ratio (Fig. 1b) and in time spent with a social target (Fig. 1c,d) compared to stress-naïve mice. A single subanesthetic, intraperitoneal (ip) injection of ketamine (10 mg/kg) administered 24 hours before the test for each behavioral assay, rescued these hedonic deficits (Fig. 1b-d). 24 hours will be used for all experiments in the paper because it captures the sustained behavioral efficacy of ketamine which is known to outlast drug clearance^10, 22^.

**Fig. 1.**
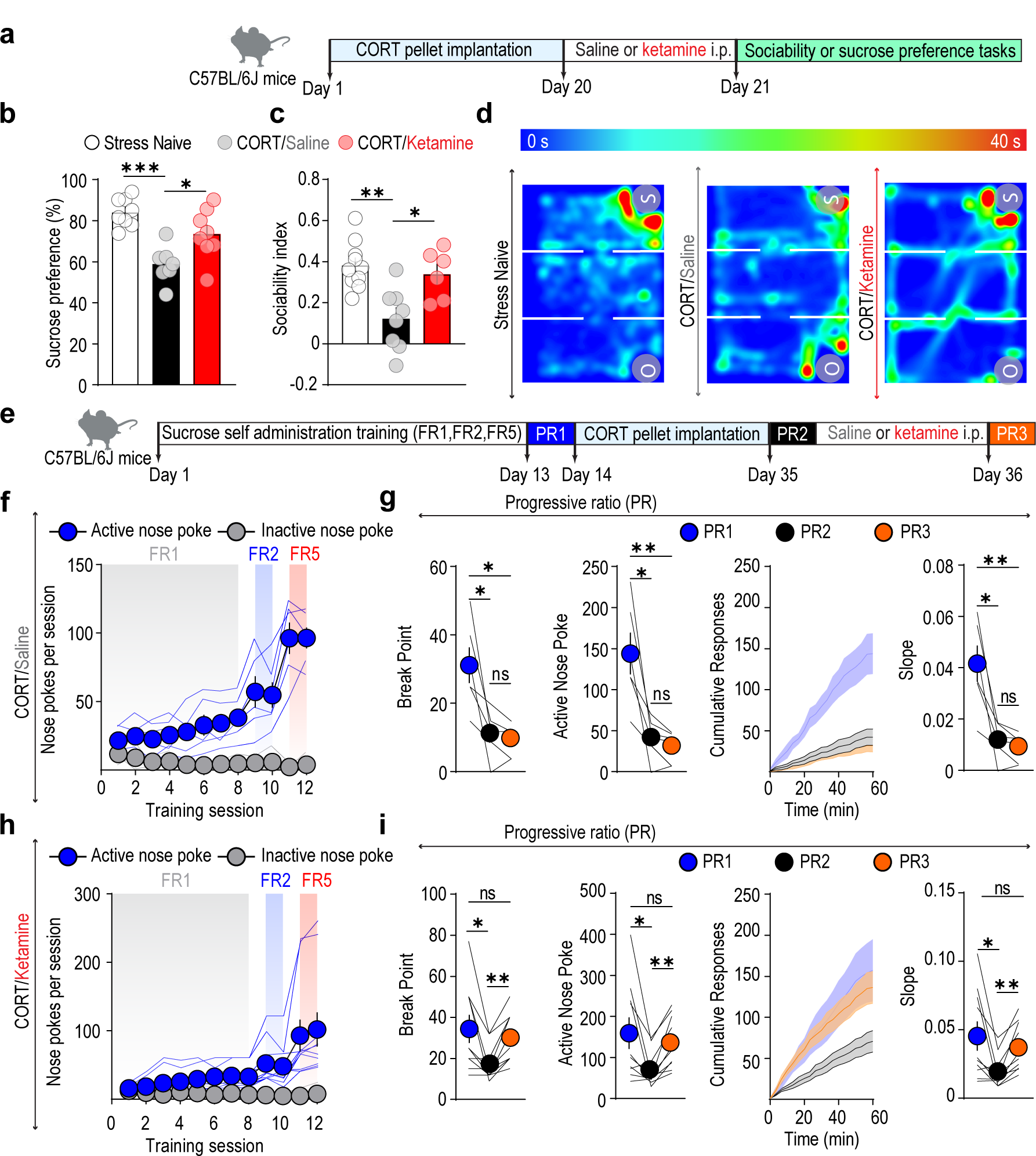
Ketamine rescues stress-induced anhedonia. **a**, **e,** Experimental timelines. **b**, Sucrose preference in stress-naïve mice (*n* = 8) and stressed mice that received saline (*n* = 7) or ketamine (*n* = 8) injections. **c**, Sociability index in stress-naïve mice (*n* = 10) and stressed mice that received saline (*n* = 8) or ketamine (*n* = 6) injections. **d**, Representative occupancy plots from a stress-naïve mouse (left) and stressed mice treated with saline (center) or ketamine (right). **f**, **h,** Performance during acquisition of the sucrose self-administration in stressed saline-(**f**, *n* = 6) or ketamine-treated (**h**, *n* = 10) mice. **g**, **i,** Break points (left), total responses (center left), cumulative responses (center right), and slopes of cumulative responses (right) during the progressive ratio tests from stressed mice who received saline (**g**, *n* = 6) or ketamine (**i**, *n* = 10) injections. Data are mean ± s.e.m. ns, not significant. \**P* < 0.05, \*\**P* < 0.01, \*\*\**P* < 0.001.

To assess the effects of chronic stress on effort-related motivation, we employed a progressive ratio (PR) schedule of reinforcement for sucrose rewards that allowed a longitudinal analysis of motivation within the same subjects at each stage in the treatment process. Mice were subjected to a PR procedure before CORT exposure (PR1), after chronic stress (PR2) and then again 24 hours after a single injection of ketamine or saline (PR3) (Fig. 1e). We found that mice exhibited decreased motivation to work for sucrose rewards after exposure to CORT while a single subanesthetic ip injection of ketamine rescued this deficit (Fig. 1f-i). Importantly, mice implanted with placebo pellets did not show any motivational impairment when tested in the PR procedure before and after placebo exposure (Extended Data Fig. 1 a,b). Taken together, these data suggest that ketamine induces a strong and sustained behavioral amelioration in several components of stress-induced anhedonia in mice, including sensitivity to reward, social motivational processes as well as effort-related motivation.

## Increased AMPAR-mediated synaptic transmission on D1-MSNs is necessary to drive the anti-anhedonic effects of ketamine

The Nucleus Accumbens (NAc) plays a major role in the generation of motivated behaviors and its structure and function are sensitive to stress^25, 32–36^. The NAc primarily contains GABAergic medium spiny neurons (MSNs) that are dichotomous in their predominant expression of either D1 or D2 dopamine receptors (D1-MSNs and D2-MSNs, respectively)^37, 38^. The respective roles of these genetically identified MSN subtypes in regulating stress-mediated behaviors differ profoundly^25, 39, 40^. Here we explored whether a single *in vivo* exposure to ketamine rescues CORT-induced anhedonia by modifying synaptic function in the NAc, accounting for potential MSN subtype differences.

Mice expressing fluorescent proteins exclusively in D1-MSNs (tdTomato line) or in D2-MSNs (A2A-cre x A14 ROSA26-tdTomato line) were subjected to chronic CORT treatment, followed by a single ip injection of either saline or ketamine (Fig. 2a, Extended Data Fig. 2a). After 24h, mice were sacrificed and acute slices containing the NAc were dissected for *ex vivo* whole-cell recordings. Stress-naïve littermates of the same genotype were used as controls. We performed targeted recordings from tdTomato positive cells, and isolated AMPAR- and NMDAR-mediated spontaneous excitatory postsynaptic currents (sEPSCs). CORT-mice treated with saline displayed a decrease in amplitude and frequency of AMPAR sEPSCs selectively in D1-MSNs compared to stress-naïve subjects (Fig. 2b and Extended Data Fig. 2b). Strikingly, a single exposure to ketamine rescued this CORT-induced reduction in amplitude and frequency of sEPSCs (Fig. 2b). Moreover, this ketamine-induced increase in AMPAR-mediated synaptic transmission was paralleled by an increase in AMPAR/NMDAR ratio, a measure of synaptic plasticity, selectively in D1-MSNs (Fig. 2c, Extended Data Fig. 2c). The observed synaptic adaptation was selective for AMPAR-mediated synaptic transmission since neither NMDAR currents nor the decay time of their kinetics were affected by ketamine administration compared to saline-treated or stress-naïve subjects in either D1- or D2-MSNs (Extended Data Fig. 3).

**Fig. 2.**
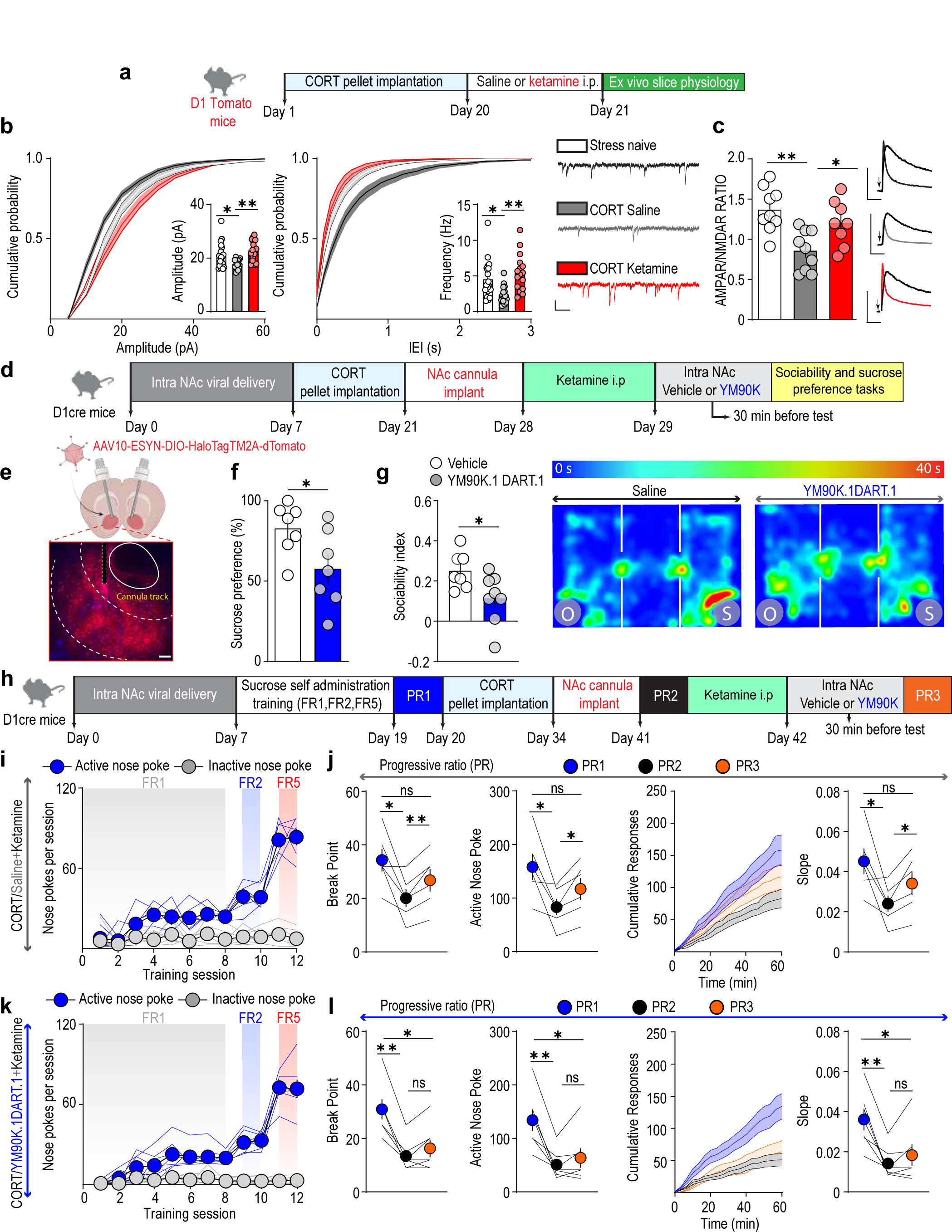
Increased AMPAR-mediated synaptic transmission on D1-MSNs is necessary to drive the anti-anhedonic effects of ketamine. **a**, **d**, **h,** Experimental timelines. **b,** Cumulative probability plots of the amplitudes (left) or frequencies (center) of sEPSCs recorded from NAc D1-MSNs in brain slices from stress-naïve mice (*n* = 20 cells; 10 mice) and stressed mice treated with saline (*n* = 15 cells; 8 mice) or ketamine (*n* = 16 cells; 9 mice). Insets: histograms of the means obtained from sEPSC amplitude (left) or frequency (center). Right, example sEPSC traces. Scale bar, 100 ms, 20 pA. **c**, Left, AMPAR/NMDAR ratio in NAc D1-MSNs from stress-naïve mice (*n* = 9 cells; 5 mice) and stressed mice who received saline (*n* = 9 cells; 5 mice) or ketamine (*n* = 8 cells; 4 mice) injections. Right, example traces for AMPAR/NMDAR ratio. Scale bars, 50 ms, 100 pA. **e**, Representative brain coronal section showing AAV10-ESYNDIO-HaloTagTM2A-dTomato transduction and cannula implantation track in the NAc from a D1-cre mouse. Scale bar, 100 µm. **f**, Sucrose preference in stressed ketamine-treated mice that received vehicle (*n* = 7) or YM90K.1^-DART.1^ (*n* = 7) micro-injections in the NAc. **g**, Left, sociability index in stressed, ketamine-treated mice that received vehicle (*n* = 7) or YM90K.1^-DART.1^ (*n* = 8) micro-injections in the NAc. Right, representative occupancy plots from stressed ketamine-treated mice who received vehicle or YM90K.1^-DART.1^ micro-injections in the NAc. **i**, **k**, Performance during acquisition of the sucrose self-administration in stressed, ketamine-treated mice that received vehicle (**i**, *n* = 6) or YM90K.1^-DART.1^ in the NAc (**k**, *n* = 7). **j**, **l**, Break points (left), total responses (center left), cumulative responses (center right), and slopes of cumulative responses (right) during the progressive ratio tests from stressed ketamine-treated mice that received vehicle or YM90K.1^-DART.1^ micro-injections in the NAc. Data are mean ± s.e.m. ns, not significant. \**P* < 0.05, \*\**P* < 0.01.

Collectively, these electrophysiological results suggest that *in vivo* exposure to ketamine increases AMPAR-mediated synaptic efficacy in NAc D1-MSNs. To test whether this ketamine-induced plasticity subserves specific behavioral functions, we tested for a causal link between the increase in AMPAR-mediated synaptic transmission in NAc D1-MSNs and the previously observed ketamine-induced behavioral amelioration of hedonic deficits. To do this, we selectively blocked ketamine-triggered sustained plasticity on AMPARs within genetically specified cells (D1-MSNs) by using DART (Drug Acutely Restricted by Tethering)^41^, a method that makes it possible to deliver traditional pharmaceuticals to genetically defined cells (Fig. 2d,e and Extended Data Fig. 4). In our study, the DART technology allowed for covalent capture of the DART-conjugated AMPAR antagonist YM90K.1^-DART^ to the surface of D1-MSNs. Cell-type specificity of YM90K.1^-DART^ capture was produced via viral-mediated expression of the bacterial protein HaloTag (DIO::HaloTag injected into the NAc of D1-cre mice), which sequesters the DART ligand and thus tethers the drug. The concentration of the DART antagonist for the *in vivo* experiments was determined by *ex vivo* acute slice preparation patch clamp experiments. Bath application of YM90K.1^-DART.1^ produced a dose-dependent effect on pharmacologically isolated AMPAR-mediated synaptic transmission solely in HaloTag positive cells (HT+) relative to HaloTag negative cells (HT-) (Extended Data Fig. 4c,d). Specifically, YM90K.1^-DART.1^ displayed a robust, specific, and sustained AMPAR-mediated inhibition at 3 µM while losing specificity at higher concentrations (Extended Data Fig. 4c,d).

**Fig. 3.**
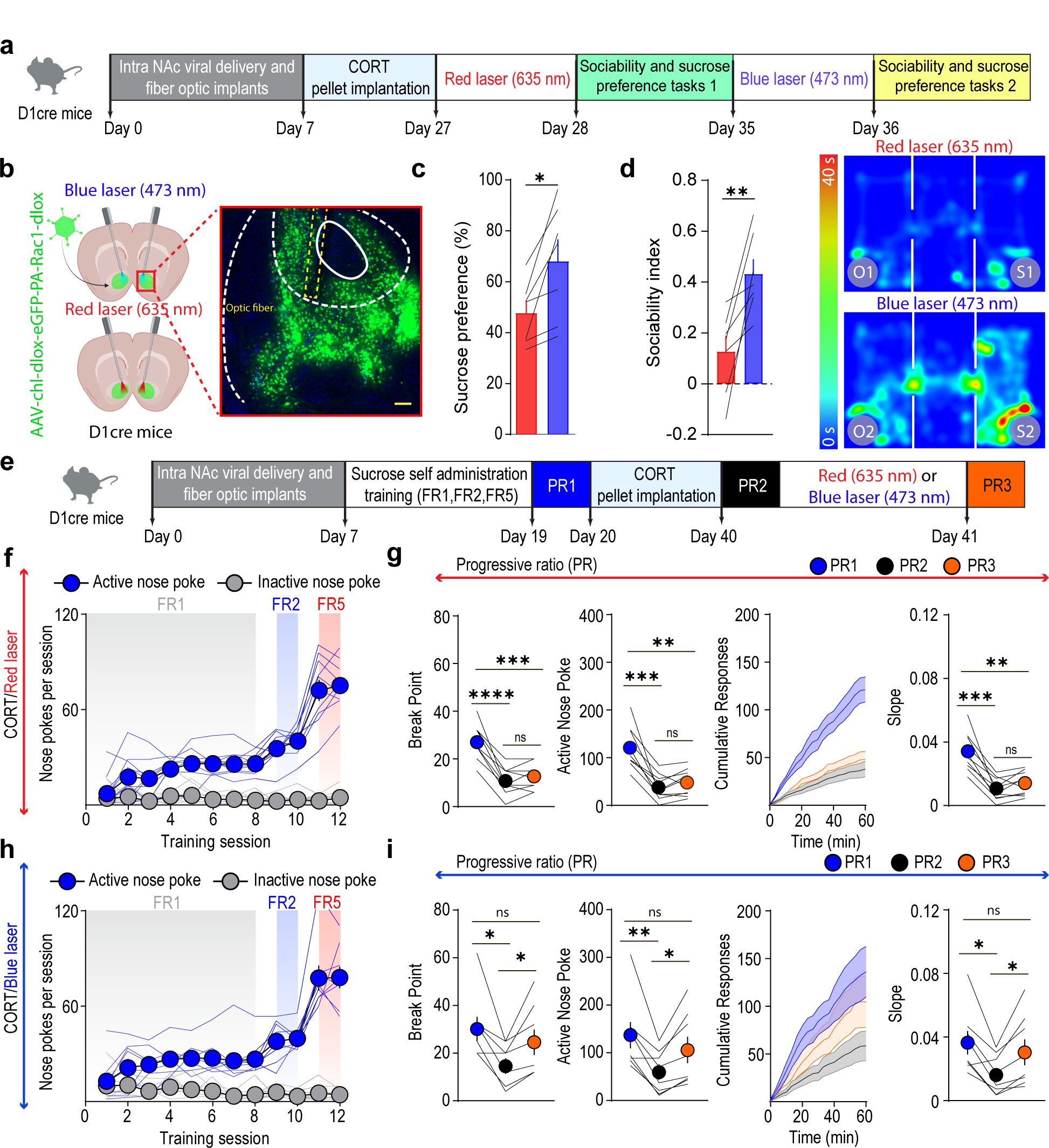
Increased AMPAR-mediated synaptic transmission on D1-MSNs is sufficient to drive anti-anhedonic effects. **a, e** Experimental timelines. **b**, Left, schematic of the experiment. Right, representative brain coronal section showing AAV-chl-dlox-eGFP-PA-Rac1-dlow transduction and fiber optic implantation track in the NAc from a D1-cre mouse. Scale bar, 100 µm. **c**, Sucrose preference in stressed mice that received red or blue stimulation in the NAc (*n* = 7). **d**, Left, sociability index in stressed mice that received red or blue laser stimulation in the NAc (*n* = 7). Right, representative occupancy plots in stressed mice that received red or blue laser stimulation in the NAc **f**, **h**, Performance during acquisition of the sucrose self-administration task in stressed mice that received red (**f**, *n* = 10) or blue laser (**h**, *n* = 8) stimulation in the NAc. **g**, **i**, Break points (left), total responses (center left), cumulative responses (center right), and slopes of cumulative responses (right) during the progressive ratio tests from stressed mice that received red (**g**, *n* = 10) or blue (**i**, *n* = 8) laser stimulation in the NAc. Data are mean ± s.e.m. ns, not significant. \**P* < 0.05, \*\**P* < 0.01, \*\*\**P* < 0.001, \*\*\*\**P* < 0.0001.

Given the specificity and robustness of YM90K.1^-DART.1^ at 3 µM, we tested this concentration *in vivo*. D1-cre mice received bilateral microinjections of AAV-DIO-HaloTag virus and cannula implantation within the NAc (Fig. 2d). Subsequently, mice were implanted with CORT-releasing pellets and then injected with ketamine 21 days later. 24 hours after ketamine administration, mice received an intra-NAc microinjection of either vehicle or YM90K.1^-DART.1^ and were tested in the sociability and sucrose preference assays beginning 30 min later. Strikingly, mice exposed to the YM90K.1^-DART.1^ microinjection failed to display a robust amelioration in sucrose preference ratio and sociability index induced by ketamine compared to mice exposed to vehicle (Fig. 2f,g).

In another set of experiments, we used the same experimental approach and applied it to the sucrose self-administration procedure (Fig. 2h). Specifically, after recovering from stereotaxic surgery, mice subjected to the sucrose self-administration procedure were tested in a PR schedule of reinforcement (PR1). Mice were retested after chronic CORT administration (PR2), then treated with systemic ketamine and tested again after receiving either vehicle or YM90K.1^-DART.1^ microinjections within the NAc. Notably, mice who received YM90K.1^-DART.1^ microinjection did not display significant amelioration in the progressive ratio task while those who received vehicle did (Fig. 2i-l). Importantly, YM90K.1^-DART.1^ at a concentration of 3 µM did not alter velocity or total distance travelled in stress-naïve mice (Extended Data Fig. 4f). By using DART.2 which improves the efficiency of chemical capture and enables cell-specific accumulation of drug to ∼3,000-times^42^, YM90K.1^-DART.2^ at 3 µM did not affect sucrose preference and sociability, while a higher concentration of YM90K.1^-DART.2^ (30 µM) reduced preference for sucrose or time spent with a conspecific in stress-naïve mice (Extended Data Fig. 4i,j). Collectively, these data suggest that AMPAR-mediated enhancement in synaptic strength within NAc D1-MSNs is necessary for ketamine-mediated amelioration of stress-induced anhedonia.

## Increased AMPAR-mediated synaptic transmission on D1-MSNs is sufficient to drive anti-anhedonic effects

We next examined whether increased AMPAR-mediated plasticity onto D1-MSNs is sufficient for ameliorating stress-induced anhedonia. To this aim, we performed a ketamine-free experiment in which we increased AMPAR-mediated synaptic transmission *in vivo* in a cell-type specific (D1-MSNs) and spatiotemporal manner. To test this, we delivered a photoactivatable (PA) version of Rac1(AAV8.2-hSyn1-chl-dlox-eGFP-PA-Rac1-(Rev)-dlox-WPRE-bGHp(A)) within the NAc (Fig. 3a,b and Extended Data Fig. 5a)^43^. PA-Rac1 is a modified Rac1, a small guanosine triphosphatase (GTPase) that enables the recruitment and clustering of AMPARs to synapses when selectively activated by blue light (473 nm)^44–46^. Therefore, this approach allows to enhance excitatory synaptic transmission by potentiating AMPAR function with spatiotemporal and cell-type specificity.

First, we confirmed that blue light exposure (473 nm) onto PA-RAc1 expressing D1-MSNs triggered a long-lasting potentiation of AMPAR currents (Extended Data Fig. 5b). In contrast, red light exposure (635 nm) did not affect AMPAR-mediated synaptic transmission and as such was implemented as a control manipulation for the *in vivo* optogenetic experiments (Extended Data Fig. 5b)^47^. To allow photoactivation of Rac1 *in vivo*, we delivered PA-Rac1 bilaterally into the NAc of D1-cre mice together with optic fibers above the injection sites (Fig. 3b). Mice were implanted with CORT-releasing pellets 7 days after surgery. 14 days after the start of chronic CORT administration, mice were exposed for 1 hour to red laser (inert manipulation), and 24 hours later they were subjected to sucrose preference and sociability assays to assess anhedonic behaviors. One week later, mice were exposed for 1 hour to blue laser stimulation and 24 hours later they were subjected to the same behavioral assays. Strikingly, mice exposed to blue laser stimulation displayed robust amelioration of hedonic deficits compared to the behavioral performances displayed after red-laser stimulation in control mice (Fig. 3c,d).

In another set of experiments, we used the same experimental approach and applied it to the sucrose self-administration procedure (Fig. 3e). Specifically, after recovering from stereotaxic surgery for viral delivery and optic fiber implantation, mice were subjected to sucrose self-administration procedure and tested in a PR schedule of reinforcement (PR1). Mice were retested after chronic CORT administration (PR2), treated with red or blue lasers, and tested again 24 hours later (PR3). Notably, mice that received blue laser stimulation displayed a significant amelioration in the progressive ratio task whereas mice exposed to red laser did not (Fig. 3f-i). Collectively, these experiments suggest that temporal-, region- and cell type-specific augmentation of AMPAR-mediated transmission within NAc D1-MSNs is sufficient to rescue CORT-induced hedonic deficits.

## Ketamine rescues stress-induced synaptic deficits within D1-MSNs at specific monosynaptic inputs to the NAc

Thus far, we have established that AMPAR-mediated synaptic transmission within NAc D1-MSNs is necessary and sufficient for amelioration of stress-induced anhedonia. To expand this mechanistic insight toward a circuit-level understanding, we sought to ask whether ketamine-induced potentiation of AMPAR-mediated synaptic transmission onto D1-MSNs occurs at specific monosynaptic inputs to the NAc. We expressed channelrhodopsin in the main glutamatergic inputs to the NAc, including the medial prefrontal cortex (mPFC), basolateral amygdala (BLA), ventral hippocampus (VH) and paraventricular nucleus of the thalamus (PVT) of D1tomato mice (Fig. 4a,b and Extended Data Fig. 6)^48^. After recovering from surgery, mice underwent chronic CORT administration and then were exposed to a systemic injection of either saline or ketamine. After 24hr, whole-cell recordings from *ex vivo* acute slices of the NAc were performed while optogenetically activating individual monosynaptic inputs to NAc D1-MSNs (Fig. 4a,b). Light-evoked AMPAR-EPSCs from optogenetically delineated inputs were normalized to NMDAR-mediated currents (AMPAR/NMDAR ratio). We found that mPFC and VH pathways to NAc D1-MSNs undergo weakening in synaptic strength after chronic CORT exposure, suggesting these monosynaptic inputs display heightened sensitivity to stress (Fig. 4b). Importantly, both mPFC and VH synapses to NAc D1-MSNs also showed enhancement in synaptic strength 24 h after *in vivo* exposure to a single subanesthetic dose of ketamine compared to saline-treated mice (Fig. 4b). In contrast, PVT and BLA synapses to NAc D1-MSNs did not show any change in synaptic strength in mice exposed to CORT and treated with either saline or ketamine compared to stress-naïve control mice (Fig. 4b). Collectively, these experiments suggest that CORT-induced reduction in amplitude and frequency of sEPSCs may arise from selective disruption in information flow provided from discrete monosynaptic inputs to the NAc while ketamine rescues these changes in synaptic efficacy.

**Fig. 4.**
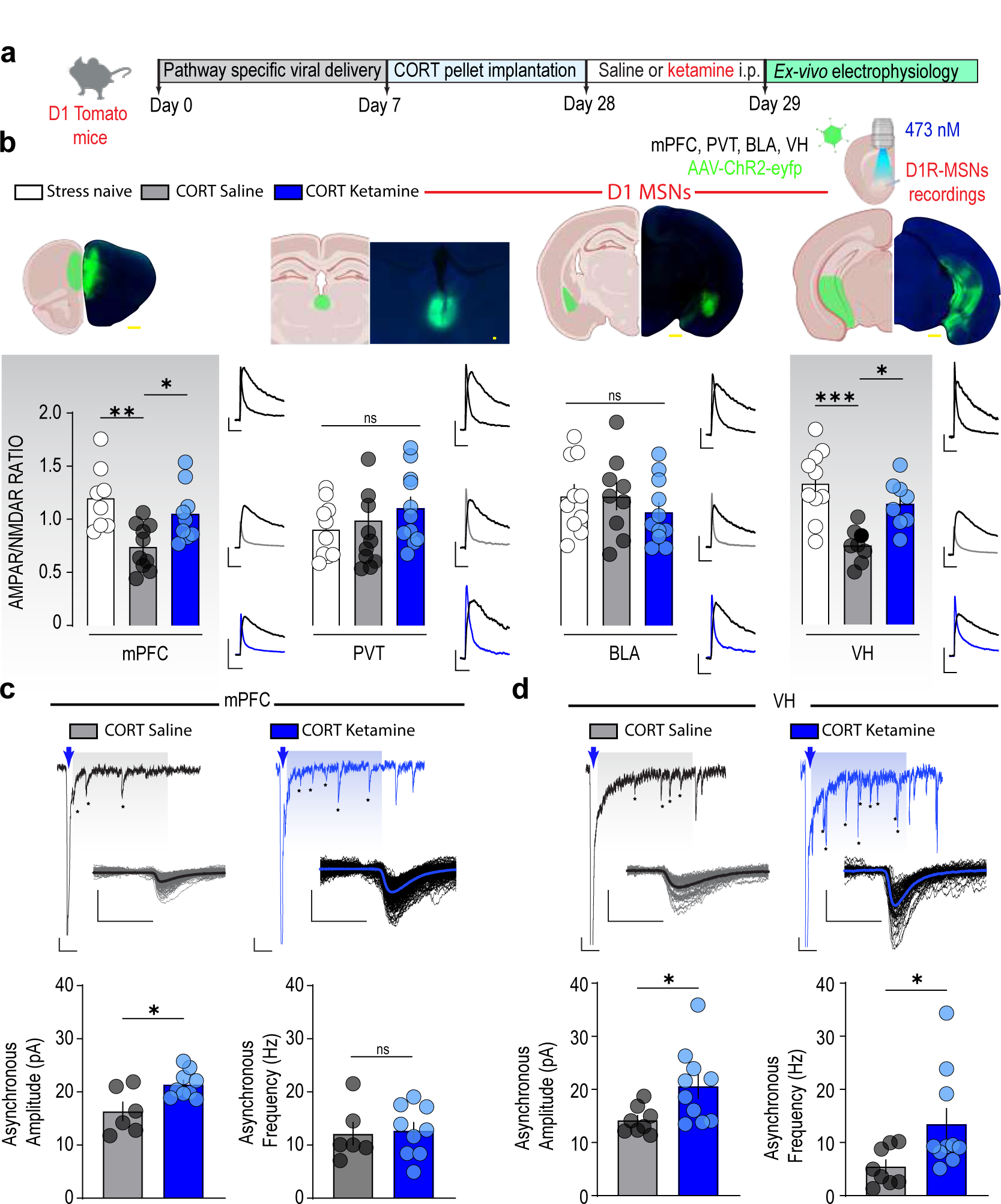
Ketamine rescues stress-induced synaptic deficits within D1-MSNs at specific monosynaptic inputs to the NAc. **a,** Experimental timeline. **b**, Top, representative brain coronal sections showing AAV-CamKIIa-hChR2(H134R)-eyfp transduction in monosynaptic inputs to the NAc (mPFC, PVT, BLA and VH). Scale bar, 500 µm. Bottom, quantification of AMPAR/NMDAR ratio in NAc D1-MSNs evoked at mPFC, PVT, BLA and VH inputs from stress-naïve mice (mPFC: *n* = 8 cells, 5 mice; PVT: *n* = 10 cells, 5 mice; BLA: *n* = 10 cells, 6 mice; VH: *n* = 10 cells, 6 mice) and stressed mice that were treated with saline (mPFC: *n* = 10 cells, 6 mice; PVT: *n* = 11 cells, 5 mice; BLA: *n* = 9 cells, 6 mice; VH: *n* = 8 cells, 5 mice) or ketamine (mPFC: *n* = 9 cells, 6 mice; PVT: *n* = 11 cells, 5 mice; BLA: *n* = 11 cells, 6 mice; VH: *n* = 8 cells, 5 mice). Example traces for AMPAR/NMDAR ratio from each input. Scale bars, 100 ms, 100 pA. **c**, **d**, Top, light-evoked mPFC (**c**) and VH (**d**) qEPSCs at D1-MSNs recorded in the presence of strontium. Blue arrow indicates light pulse. Asterisks indicate qEPSCs. Scale bars, 100 ms, 20 pA. Insets: Overlay of individual (grey and black) and average (black and blue) onset-aligned qEPSCs from examples. Scale bars, 10 ms, 20 pA. Bottom, summary of the amplitude and frequency of mPFC (**c**) and VH (**d**) qEPSCs at D1-MSNs from stressed mice that received saline (mPFC: *n* = 6 cells, 4 mice; VH: *n* = 8 cells, 5 mice) or ketamine (mPFC: *n* = 9 cells, 5 mice; VH: *n* = 10 cells, 6 mice). Data are mean ± s.e.m. ns, not significant. \**P* < 0.05, \*\**P* < 0.01, \*\*\**P* < 0.001.

To further test the hypothesis that ketamine-triggered AMPAR-mediated synaptic plasticity occurs at specific monosynaptic inputs to accumbal D1-MSNs, we analyzed quantal release events by replacing calcium with strontium (Sr^2+^) in the extracellular solution following either optogenetically-evoked mPFC or VH afferent stimulation (Fig. 4c,d)^49^. The resultant input-specific quantal EPSCs (qEPSCs) provide a direct and quantitative measure of unitary synaptic strength (amplitude) as well as an indirect estimate of the number of connections (frequency). Optogenetic stimulation of mPFC axons evoked asynchronous release displaying equal frequency of qEPSCs in saline- and ketamine-treated subjects (Fig. 4c). In contrast, qEPSC amplitude was larger in the ketamine-treated group (Fig. 4c). Therefore, these results suggest that the locus of expression of synaptic efficacy at mPFC to NAc D1-MSNs is postsynaptic. Conversely, light-induced stimulation of VH axons evoked asynchronous release displaying both higher frequency and larger amplitude of qEPSCs in ketamine-treated group, suggesting changes in postsynaptic efficacy and possibly a larger number of connections (Fig. 4d). Taken together, these findings suggest that ketamine-evoked synaptic plasticity occurs within NAc D1-MSNs by rescuing synaptic efficacy at discrete monosynaptic inputs to the NAc (mPFC and VH) and that the underlying mechanisms may be unique to each pathway.

## Input-specific control of ketamine-mediated rescue of stress-induced anhedonia

Finally, we wanted to dissect NAc input-specific regulation of ketamine’s rescue of stress-induced anhedonia. Thus, we inhibited information flow in each specific pathway chemogenetically and assessed whether mPFC or VH activity is required for ketamine’s amelioration of CORT-induced anhedonia. To do this, we employed an intersectional genetic approach, injecting a retrograde cre-dependent virus bilaterally within the NAc, and either AAV-DIO-eYFP (control) or AAV-DIO-HA-hM4D(Gi)-IRES-mCitrine (experimental group) within either the mPFC or VH (Fig. 5a and Extended Data Fig. 7)^50^. After recovery from surgery, we administered CNO (5 mg/kg) or vehicle ip 30 min prior to ketamine injection in CORT-treated mice. 24 hour later, we tested mice in the sucrose preference, sociability and sucrose self-administration assays. We found that mice expressing hM4D but pretreated with vehicle displayed robust amelioration of sucrose preference ratio and sociability index, as well as in the number of rewards obtained in the sucrose self-administration procedure during the progressive ratio regimen (Fig. 5b-f and Extended Data Fig. 7). In contrast, inhibiting either mPFC or VH activity to the NAc prior to ketamine delivery was sufficient to abolish rescue from CORT-induced anhedonia in all three behavioral assays (Fig. 5b-f). Intriguingly, preventing information flow to the NAc prior to ketamine exposure led to a different pattern of behavior for mPFC- and VH-pathway silencing. Specifically, inhibition of the VH pathway resulted in mice having an increased latency to first entry into the social chamber and to first nose poke for a sucrose reward, and it also affected the total number of social entries (Fig. 5b-f). On the contrary, inhibition of the mPFC pathway to the NAc did not affect how ketamine rescues initiation of hedonic responses (Fig. 5b-f). Importantly, CNO injections in mice expressing AAV-DIO-eYFP in VH and mPFC did not affect behavioral performance in sucrose preference or sociability tasks (Extended Data Fig. 7e,f). Collectively, these data suggest that both mPFC- and VH-NAc pathways are crucial for ketamine’s anti-anhedonic effects, however they exert different control over various components of hedonic behaviors. Specifically, VH-NAc synapses are uniquely poised to control the initiation to work for obtaining a reinforcer as well as the initiation of both approach and return to social encounters, whereas mPFC-NAc synapses play a more general role in sustaining task engagement.

**Fig. 5.**
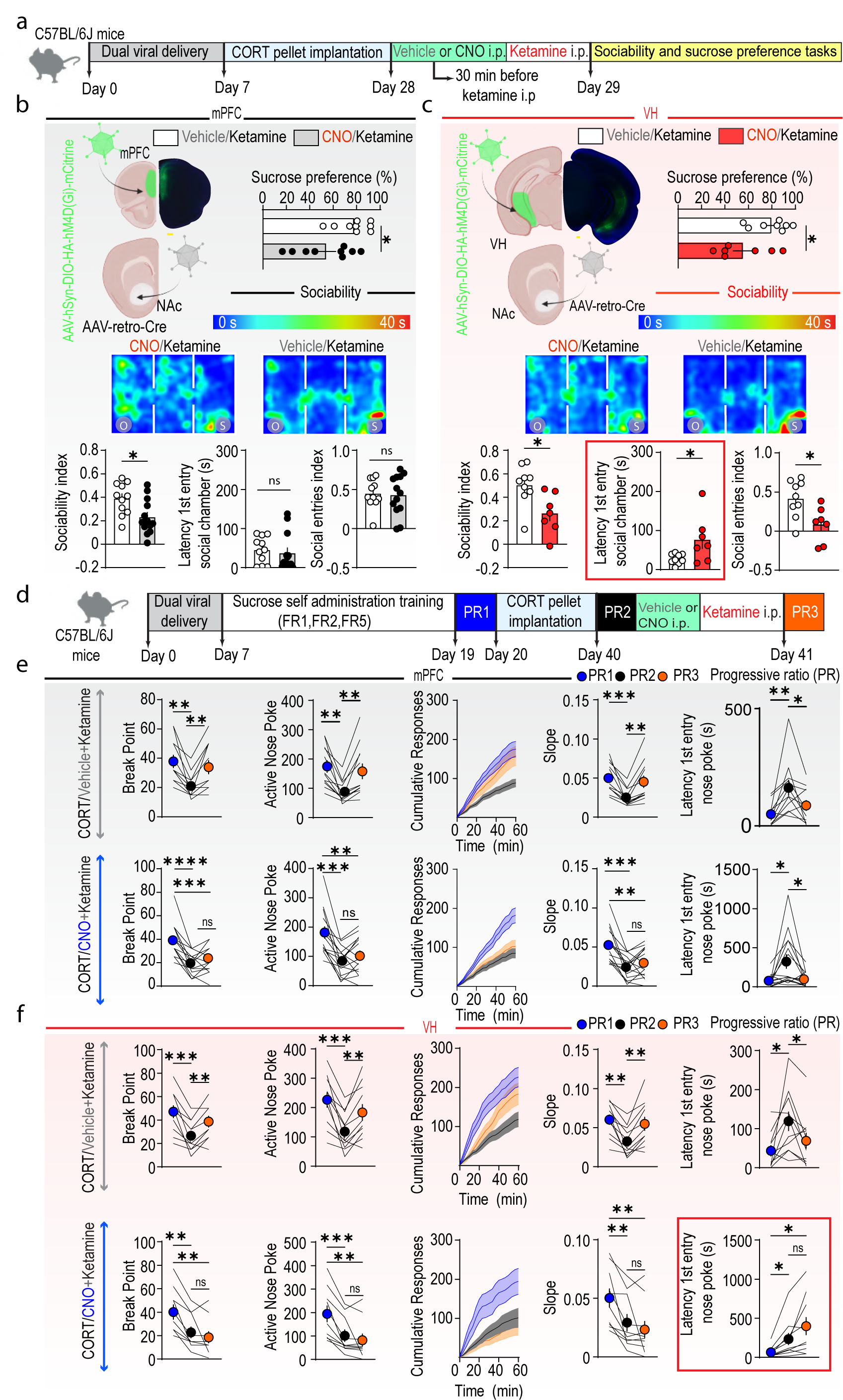
Input-specific control of ketamine-mediated rescue of stress-induced anhedonia. **a, d,** Experimental timelines. **b, c,** Top left, representative brain coronal sections showing AAV-hSyn-DIO-HA-hM4D(Gi)-IRES-mcitrine transduction in mPFC (**b**) or VH (**c**) to NAc projecting neurons. Scale bar, 500 µm. Top right, sucrose preference of stressed mice that received vehicle or CNO injections before ketamine treatment in the mPFC (**b**, vehicle: *n* = 9, CNO: *n* = 9) or VH (**c**, vehicle: *n* = 8, CNO: *n* = 7) groups. Center, representative occupancy plots from stressed mice that received vehicle (right) or CNO (left) before ketamine treatment in the mPFC (**b**) or VH (**c**) groups. Bottom, sociability index, latency to first entry in the social chamber and social entries index in stressed mice who received vehicle or CNO injections before ketamine treatment in the mPFC (**b**, vehicle: *n* = 11, CNO: *n* = 13) or VH (**c**, vehicle: *n* = 9, CNO: *n* = 7) groups. **e, f,** Break points (left), total responses (center left), cumulative responses (center), slopes of cumulative responses (center right), and latency to first nose-poke (right) during the progressive ratio tests from stressed mice that received vehicle or CNO injections before ketamine treatment in the mPFC (**g**, vehicle: *n* = 12, CNO: n = 15) or VH (**h,** vehicle: *n* = 11, CNO: n = 10) groups. Data are mean ± s.e.m. ns, not significant. \**P* < 0.05, \*\**P* < 0.01, \*\*\**P* < 0.001, \*\*\*\**P* < 0.0001.

## Concluding remarks

Our findings elucidate a novel framework for the role of ketamine in rescuing stress-induced anhedonia via synaptic action within NAc D1-MSNs. By combining cell-type specific pharmacology, synaptic electrophysiology and *in vitro* and *in vivo* opto- and chemo-genetic manipulations, we provided evidence that temporally restricted upregulation of AMPAR function selectively within NAc D1-MSNs is necessary for ketamine-elicited rescue of stress-induced anhedonia. Artificially mimicking this ketamine-induced increase in AMPAR function within D1-MSNs recapitulates the behavioral amelioration induced by ketamine. We also found that ketamine rescued stress-induced anhedonia by restoring synaptic efficacy at both mPFC- and VH-NAc D1-MSN synapses. Intriguingly, we showed an input-specific variation in control over different components of hedonic behaviors. Specifically, VH-NAc synapses control the initiation to work for obtaining a reinforcer as well as the initiation of both approach and return to social encounters, whereas mPFC-NAc synapses play a more general role in sustaining task engagement. A functional interpretation of the interplay between the roles of these two pathways in regulating anhedonia and the antidepressant effect of ketamine may depend on the ability of hippocampal afferents to gate information throughput to the NAc^51, 52^. Indeed, hippocampal input is crucial for driving NAc activity^53^, and activation of this pathway enables the NAc to modulate information throughput from the mPFC to downstream NAc targets. In line with our findings, ketamine restores lost spines and rescues coordinated activity in the mPFC in mice exposed to chronic CORT regimen^28^. In humans, functional imaging studies have discovered a ketamine-mediated increased functional connectivity within fronto-striatal neural circuitry as a neural correlate to successful treatment for depression^54–56^. Additionally, magnetic resonance imaging studies showed that ketamine exerts enhanced antidepressant effects in patients with relatively smaller hippocampus^57^, supporting the notion that ketamine’s effects may depend on a particular balance between mPFC- and hippocampal-NAc inputs. Together with our findings, these studies suggest that functional connectivity and morphometric data analyses may be used as a biomarker of antidepressant treatment outcome. Moreover, our study further highlights the importance of the reward circuitry in the establishment of both rapid and sustained therapeutic effect by ketamine exposure in the context of anhedonia^58, 59^.

In summary, our data indicate that ketamine improves appetitive motivational processes including approach behavior, exertion of effort, and task engagement by restoring proper functioning of the NAc D1-MSNs and selected glutamatergic afferents. This understanding not only improves our knowledge of ketamine’s mechanism of action, but may also guide future research for developing, in the long run, new therapeutic approaches for targeting specified circuit and cellular elements in the context of several neuropsychiatric disorders in which anhedonia represents a core symptom. This is particularly important since ketamine widespread clinical use has been limited due to its ability to trigger psychotomimetic symptoms and addiction liability^60^. Understanding the detailed circuit and synaptic mechanisms and establishing a causal link between ketamine-evoked synaptic plasticity and specific behavioral outcomes is crucial for designing novel and safer therapeutic drugs. In fact, such approach will allow harnessing the properties of ketamine into a more specific drug therapy, with fewer unwanted side effects.

## Methods

### Mice

Adult male and female mice (7-16 weeks) were bred in-house from stock lines obtained from the Jackson Laboratory (wild type C57/BL6J: JAX#000664; Ai14, B6.Cg-Gt(ROSA)26Sortm14(CAG-tdTomato)Hze/J; Drd1a-tdTomato, B6.Cg-Tg(Drd1a-tdTomato)6Calak/J) or the MMRRC (Mutant Mouse Resource and Research Center) repository (A2a-Cre, B6.FVB(Cg)-Tg(Adora2a-cre)KG139Gsat/Mmucd; D1-cre, Tg(Drd1-cre)EY217Gsat/Mmucd). Mice were housed in temperature- and humidity-controlled facilities under a reverse 12-h light-dark cycle (light off at 07:00 a.m.). All physiology and behavior experiments were performed during the dark cycle. Mice had *ad libitum* access to food and water throughout the experiments. Mice were group housed except for those used for subcutaneous pellet implantation. All animal procedures were approved by Institutional Animal Care and Use Committee at Washington University in St. Louis and were performed in accordance with the *Guide for the Care and Use of Laboratory Animals*, as adopted by the NIH.

### Drugs

(R,S)-ketamine (10 mg/kg, Sigma-Aldrich) and Clozapine N-oxide dihydrochloride (CNO, 5 mg/kg, Tocris Bioscence) were dissolved in sterile 0.9% NaCl (saline). YM90K.1^DART.1^ and YM90K.1^DART.2^ (provided by M.R. Tadross) were administered intracranially. Both male and female experimenters administered the drugs.

### Surgeries

#### Stereotaxic surgeries

Standard stereotaxic surgeries were conducted under ketamine (100 mg/kg; ip) and xylazine (10 mg/kg; ip) anesthesia. For electrophysiology, optogenetic, chemogenetic and behavioral experiments, purified and concentrated AAVs were injected at the following coordinates: NAc (anteroposterior (AP), +1.5mm; mediolateral (ML), ±0.9mm; dorsoventral (DV), −4.5mm; 500 nL), mPFC (AP, +1.8mm; ML, ±0.4mm; DV, −1.9mm; 400 nL), VH (AP, −2.9mm; ML, ±3.1mm for males and ±3.0 mm for females; DV, −4.5mm; 300 nL), BLA (AP, −1.6mm; ML, ±3.1mm; DV, −4.9mm; 300 nL) and PVT (AP, +1.4 mm; ML, ± 1.3mm; DV, −3.6mm with 22.5 degree angle; 300 nL). Bregma and skull surface at the injection site were used as reference points. AAV10-ESYNDIO-HaloTagTM2A-dTomato and AAVrh10-ESYNDIO-TM2A-dTomato (control) were provided by M.R. Tadross, prepared by the viral vector core of Duke University. AAV8-hSyn-DIO-HA-hM4D(Gi)-IRES-mcitrine, pAAVretro-EF1a-Cre, AAV1-CamKIIa-hChR2(H134R)-eyfp were obtained from Addgene. AAV-1/2-hSyn1-chl-dlox-eGFP-PA-Rac1(rev)-dlox-WPRE-bGHp(A) was obtained from the viral vector core of University of Zurich. Injections were carried out using a 2 µl Hamilton syringe connected to FEP tubing and to 29 Ga stainless steel injectors at a rate of 0.1 µL min^-1^; injectors were left *in situ* for an additional 5 min to allow for the diffusion of virus; 21 d was the minimum viral expression time. For the DART experiments, cannulas were implanted bilaterally over the NAc (AP, +1.5mm; ML, ± 1.85mm; DV, −3.2mm; 12 degrees) and secured with skull screws and dental cement. For PA-Rac1 experiments, optic fibers were implanted bilaterally over the NAc (AP, +1.5mm; ML, ± 1.85mm; DV, −4.2mm; 12 degrees) and secured with skull screws and dental cement. All mice were left to recover for at least 1 week before starting behavioral experiments. Only mice with verified infection sites, cannula and fiber placements were included in the analyses.

#### Subcutaneous implantation of pellet

For implanting corticosterone pellets (#G-111, Innovative Research of America), mice were anaesthetized with ketamine (100 mg/kg; ip) and xylazine (10 mg/kg; ip) and a small incision was made on the back of the mouse. A pellet was positioned in the incision between the skin and the underlying muscle tissue. The skin incision was closed with monofilament nylon suture. The pellets allow for constant release of corticosterone because they are made of a biodegradable matrix of cholesterol and cellulose. We used pellets corresponding to a dose of 50 mg/kg corticosterone per day.

### Patch-clamp electrophysiology

Whole-cell patch-clamp recordings were carried out as previously described^61^. Specifically, mice were anaesthetized with isofluorane (Pivetal) before decapitation. Brains were rapidly removed and placed in ice-cold NMDG-based cutting solution (in mM: 92 NMDG, 20 HEPES, 25 glucose, 30 NaHCO_3_, 2.5 KCl, 1.2 NaPO_4_, 5 sodium ascorbate, 3 sodium pyruvate, 2 thiourea; osmolarity: 303-306 mOsm) and continuously bubbled with 95% O_2_/5% CO_2_. Coronal sections (300 µm) containing the NAc were obtained at a speed of 0.07 mm/s with a Leica VT1200 Vibratome while the brain was submerged in ice-cold NMDG-based cutting solution. Slices were subsequently incubated in a NMDG-based cutting solution for 5-10 min at 34°C. Slices were then transferred in artificial cerebrospinal fluid (aCSF) (in mM: 92 NaCl, 20 HEPES, 25 glucose, 30 NaHCO_3_, 2.5 KCl, 1.2 NaPO_4_, 5 sodium ascorbate, 3 sodium pyruvate, 2 thiourea; osmolarity: 303-306 mOsm) saturated with 95% O_2_/5% CO_2_ at room temperature. Slices were allowed to incubate in this solution for at least 1 hr before being transferred to the recording chamber. The recording chamber was kept at 32°C and perfused with a pump (World Precision Instruments) at a flow rate of 1.5-2.0 ml per minute with aCSF (in mM: 126 NaCl, 2.5 KCl, 1.4 NaH_2_PO_4_, 1.2 MgCl_2_, 2.4 CaCl_2_, 25 NaHCO_3_, and 11 glucose; osmolarity: 303-305 mOsm). Whole-cell recordings of spontaneous and evoked EPSCs were made utilizing glass microelectrodes (2-3 MΩ) filled with cesium-based internal solution (in mM: 117 cesium methanesulfonate, 20 HEPES, 0.4 EGTA, 2.8 NaCl, 5 TEA-Cl, 4 Mg-ATP, 0.4 Na-GTP, and 5 QX-314; osmolarity: 280-285 mOsm). To isolate EPSCs, cells were voltage clamped at −70 mV and 100 µM picrotoxin was included in the aCSF. For the AMPAR/NMDAR ratio experiments, AMPAR-mediated currents were isolated with the selective NMDAR antagonist AP5 (50 µM). The NMDAR-mediated current was digitally obtained by taking the difference between the current before and after AP5 bath application. For obtaining asynchronous EPSCs, extracellular calcium was substituted with Strontium (4 mM) in the superfused aCSF. Asynchronous EPSCs were analyzed during a 500 ms window beginning 5 ms after optical stimulation. Whole-cell recordings of intrinsic excitability were made utilizing glass microelectrodes (3-4 MΩ) containing (in mM): 135 K-gluconate, 10 HEPES, 4 KCl, 4 Mg-ATP, and 0.3 Na-GTP. For validation of chemogenetic experiments, 500-ms somatic depolarizing current steps were delivered up to 300 pA before and after bath application of CNO (10 µM). Medium-spiny neurons (MSNs) were identified using IR-DIC optics on an inverted Olympus BX5iWI microscope. TdTomato-positive cells were classified based on strong fluorescence, while TdTomato-negative cells lacked fluorescence but were adjacent to TdTomato-positive cells. Optogenetically-evoked synaptic responses were elicited using a 473 (blue) and 635 (red) nm lasers (1 ms pulse) directed at the brain slice (Thor Labs). Laser intensity was adjusted to evoke EPSCs at approximately half of maximal amplitude (1-10 mW). Synaptic responses were analyzed using ClampFit (Molecular Devices) and Easy Electrophysiology (Easy Electrophysiology Ltd) softwares. Neurons were voltage clamped at −70mV utilizing a Multiclamp 700B amplifier (Molecular Devices). Data were filtered at 1 kHz and digitized at 20 kHz using a 1440A Digidata Digitizer (Molecular Devices). Series-resistance (10-20 MΩ) was monitored using a −5 mV voltage step. Cells with >20% change in series resistance were discarded from further analysis.

### Serum corticosterone measurement

Blood was collected at 10:00 am from the lateral tail vein. Blood was collected into centrifugal tubes, allowed to clot and centrifuged (2000 ×g, 20 min, 4°C) to separate serum. The serum samples were stored at −20°C until hormone assay. Corticosterone concentrations were measured with an ELISA kit purchased from Crystal Chem (Catalog Number #80556).

### Behavioral assays

#### Sucrose self-administration

Mice operant-conditioning chambers (Med Associates, Fairfax, VT) were equipped with two nose poke holes on the left-hand wall 11 cm apart, a food magazine connected to a food pellet dispenser in the center of the left-hand wall and one house light positioned on the top right-hand wall. Mice were trained in 1 hour daily sessions. On fixed-ratio 1 (FR1) schedule of reinforcement, an active nose poke resulted in sucrose delivery in the food magazine (20 mg sucrose pellets, Bio-Serv) together with turning on the house light cue for 20 s. During this 20 s period, no further action had consequence (time out period). Poking in the inactive hole had no consequence. Mice were first trained on FR1 for 8 sessions. Then, mice underwent 2 sessions of FR2 and 2 sessions of FR5. Motivation was then assessed using a progressive ratio (PR) schedule of reinforcement (PR1). In the PR session, the number of nose pokes necessary to obtain sucrose followed the equation response ratio = (5 × e^(0.2 × reward number)^ – 5 rounded to the nearest integer resulting in the following PR steps: 1, 2, 6, 9, 12, 15, 20, 25, 32, 40, 50, 62, 77, 95. The following day, placebo or corticosterone pellets were implanted subcutaneously. Mice were tested on a second PR schedule of reinforcement (PR2) to assess any effect of chronic stress on motivation 21 days after pellet implantation.

For studies testing the effect of ketamine on motivated behavior, mice were injected with saline or ketamine following PR2 and then tested again 24 h later in an additional PR session (PR3).

For studies testing the necessity of NAc D1 AMPARs in ketamine’s effect on motivated behavior, one week before PR2, guide cannulas were placed bilaterally above the NAc. After PR2, mice were injected with saline or ketamine. The following day, mice were injected with vehicle or YM90K.1^DART.1^ in the NAc, and then 1 hour later, mice were tested again for an additional PR session (PR3).

For studies testing the sufficiency of NAc D1 AMPARs on motivated behavior, mice were stimulated with red (635 nm) or blue (473 nm) laser for 1 hour (2 s, every 20 s) after PR2 and tested again 24 h after optogenetic stimulation in an additional PR session (PR3).

For studies testing the necessity of specific inputs to the NAc in ketamine’s effect on motivated behavior, mice were injected with vehicle or CNO (10 mg/kg, Tocris) after PR2, and then 30 minutes later they were injected with ketamine. The following day mice were tested again for an additional PR session (PR3).

#### Three chamber sociability assay

The three-chamber sociability test was performed in rectangular Plexiglas arena (60 cm x 40cm x 20cm) divided into three chambers that are separated by removable doors situated on the walls of the center chamber. Mice were habituated to the arena with two empty enclosures placed in the two outer chambers for 5 min. Then, the test mouse was temporarily placed in the center chamber by closing the removable doors. A nonfamiliar conspecific age-matched mouse was placed into one of the enclosures while an object was placed into the other enclosure. The test mouse was kept in the center chamber for 2 min. The doors were then removed and the test mouse was allowed to freely explore the arena and the two enclosures for 20 min. The walls of the enclosures, consisting of vertical plexiglas bars, allowed auditory, olfactory, visual and tactile contact between the test mouse and the stimulus mouse or the object. The position of the stimuli was randomly assigned and counterbalanced. The location of the mouse was assayed automatically using a video tracking system (ANY-maze). Sociability index was calculated as: ((time in social side − time in object side)/(time in social side + time in empty side)). Social entries index was calculated as: ((number of entries to the social side – number of entries to the object side)/(number of entries to the social side + number of entries to the object side)).

#### Sucrose preference task

Mice were placed in standard rodent cages (36cm × 18cm × 12 cm) and given access to a bottle containing tap water and a bottle containing 1% sucrose overnight. The total amount of tap water and sucrose consumed was recorded. Sucrose preference is expressed as the amount of sucrose consumed divided by the total amount of water consumed multiplied by 100.

#### Locomotor activity

Mice were placed in an arena (45cm × 45 cm) for 20 min and locomotor activity (total distance travelled and velocity) was recorded and analyzed by ANY-maze.

### Statistics and reproducibility

All experiments were performed at least twice to avoid any unspecific day/condition effect. Mice were randomly assigned to treatment conditions by cage before surgery and behavioral experiments. During behavioral tasks experimenters were not always blind to the group affiliation (control vs experimental) given familiarity with the subjects. However, for histology and *ex vivo* electrophysiological recordings, the experimenters were blinded to the group assignment of the mice (control vs experimental). During electrophysiological data processing and analysis experimenters were blinded to the group affiliation until all data was processed and group comparisons could be made. All analyses were performed using the following softwares: Clampex, Easy Electrophysiology, ANY-maze, Excel, and R (http://www.r-project.org/). Visualizations and statistics were performed in GraphPad Prism version 9. All data are presented as mean ± SEM unless reported otherwise. Data were tested for normal distribution using the Shapiro–Wilk normality test. Normally and non-normally distributed datasets were then analyzed with parametric or non-parametric comparisons, respectively. If ANOVA yielded a significant main effect or interaction, post-hoc comparisons were conducted. All statistical tests were two-tailed. A P value of 0.05 was considered significant.

## Data availability

The data that support the findings of this study are available from the corresponding author upon request.

## Acknowledgements

We thank Kat Czerpaniak, Francesca R. Fiocchi, Adam Kepecs, Miriam Melis, Steven Mennerick, Yavin Shaham, Hugo Tejeda, and Charles Zorumski for their comments on the manuscript. This study was supported by the NIMH grant MH130610 (MP), the Taylor Family Institute for Innovative Psychiatric Research (MP), The Hope Center Pilot Grant (MP), The McDonnell Center for Systems Neuroscience Small Grant (FL) and the NARSAD Young Investigator Grant 27102 and P&S Fund (MP). Portions of the figures were created with BioRender (biorender.com).

## Author Contributions

FL and MP conceived the experiments with inputs from BCS, MRT and CAZJr. FL, MP, SL, JL, JR and LB performed the experiments and analyses. FL and MP wrote the manuscript with the help of all authors. MP supervised the study.

## Competing interests

The authors declare no competing interests.

---

**Extended Data Fig. 1.**
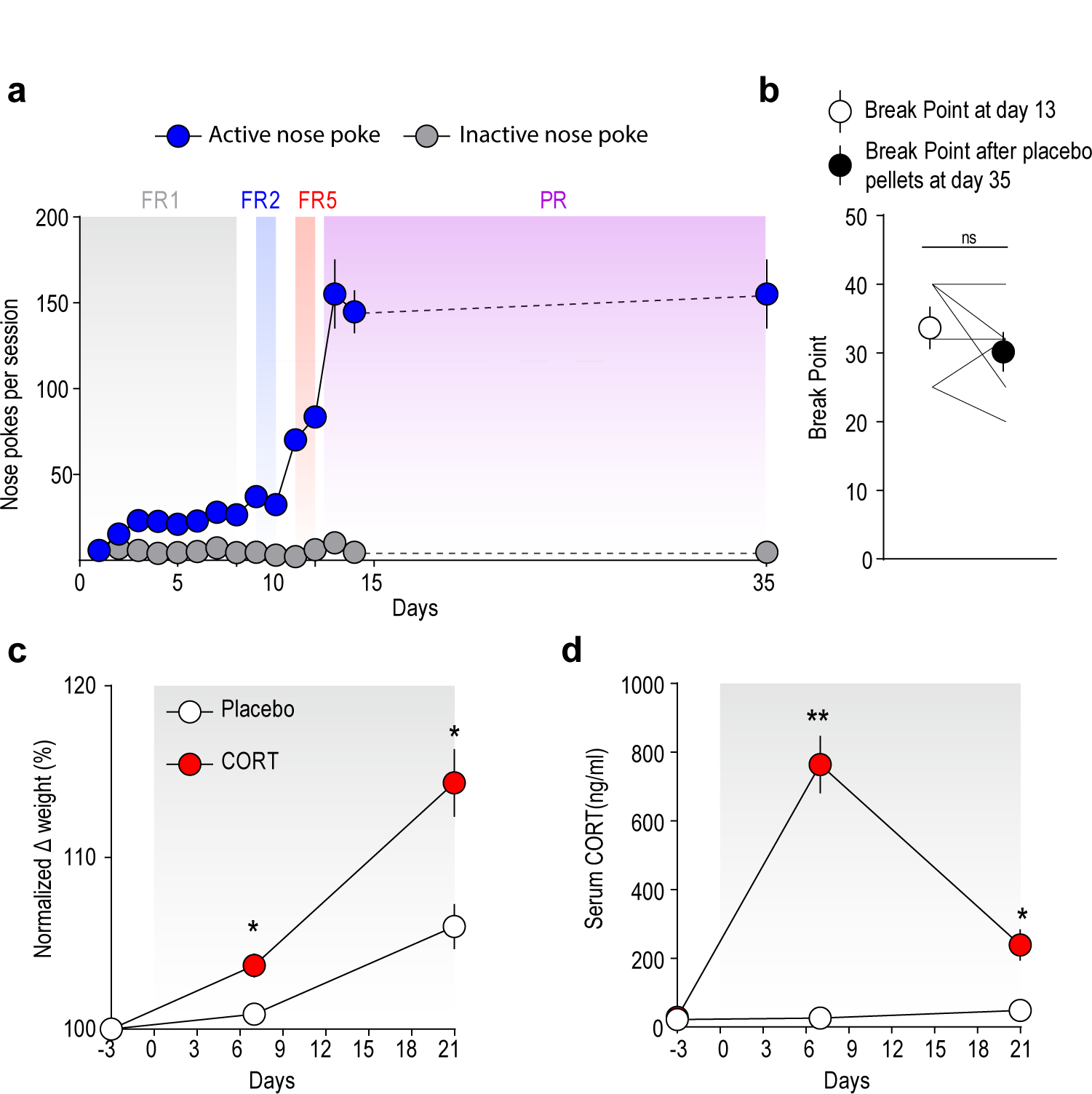
Sucrose self-administration in placebo-treated mice and corticosterone measurements in corticosterone- and placebo-treated mice. **>a,** Performance during acquisition and during the progressive ratio tests in placebo-treated mice (*n* = 6). **b,** Break points during the progressive ratio tests at days 13 and 35 from placebo-treated mice. **c, d,** Change in bodyweight (**c**) and concentration of serum corticosterone (**d**) in placebo-(n = 7) and CORT-treated mice (n = 6) at different days from pellet implantation (Day 0). Gray shading indicates post-pellet implantation period. Data are mean ± s.e.m. ns, not significant. \**P* < 0.05, \*\**P* < 0.01.

**Extended Data Fig. 2.**
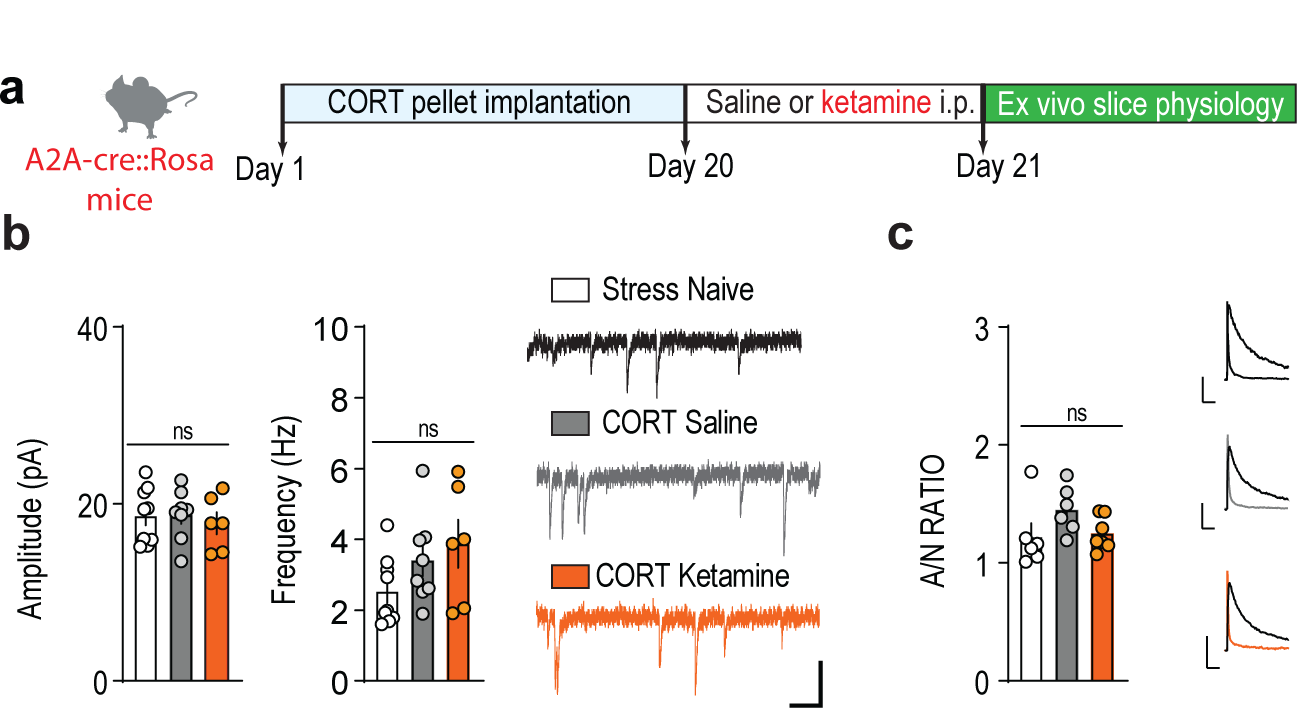
AMPAR-mediated synaptic transmission on D2-MSNs in stress-naïve mice and stressed mice that were treated with saline or ketamine. **a,** Experimental timeline. **b,** Histograms of the means obtained from sEPSC amplitude (left) or frequencies (center) recorded from D2 cells in brain slices from stress-naïve mice (*n* = 9 cells; 5 mice) and stressed mice that received saline (*n* = 8 cells; 5 mice) or ketamine (*n* = 6 cells; 4 mice). Right, example sEPSC traces. Scale bar, 100 ms, 10 pA. c, Left, AMPAR/NMDAR ratio in NAc D2-MSNs from stress-naïve mice (*n* = 6 cells; 4 mice) and stressed mice that received saline (*n* = 6 cells; 4 mice) or ketamine (*n* = 6 cells; 4 mice). Right, example traces for AMPAR/NMDAR ratio. Scale bars, 100 ms, 50 pA. Data are mean ± s.e.m. ns, not significant.

**Extended Data Fig. 3.**
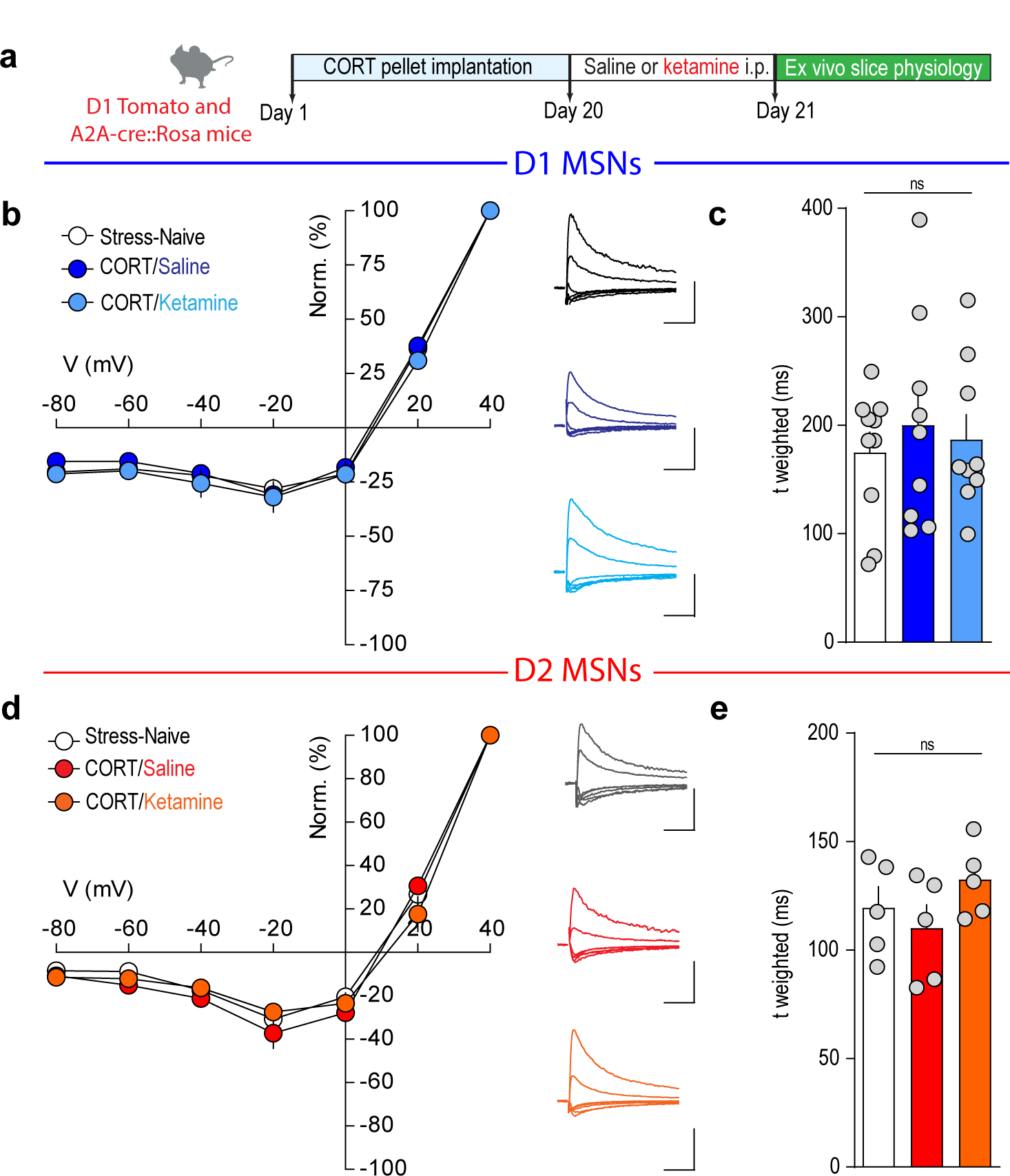
NMDAR-mediated synaptic transmission on D1- and D2-MSNs in stress-naïve mice and stressed mice that were treated with saline or ketamine. **a,** Experimental timeline. **b, d,** Right, current-voltage relationships of the electrically evoked NMDAR-mediated response recorded in D1-MSNs (**b**) and D2-MSNs (**d**) in brain slices from stress-naïve mice (D1-MSNs: *n* = 10 cells, 6 mice; D2-MSNs: *n* = 5 cells, 3 mice) and stressed mice that received saline (D1-MSNs: *n* = 9 cells, 5 mice; D2-MSNs: *n* = 5 cells, 3 mice) or ketamine (D1-MSNs: *n* = 9 cells, 5 mice; D2-MSNs: *n* = 5 cells, 3 mice). Left, example traces for NMDA I/V curves. Scale bars, 100 ms, 50 pA. c, e, Weighted decay time constants (τW) of NMDAR EPSCs recorded at +40 mV. Data are mean ± s.e.m. ns, not significant.

**Extended Data Fig. 4.**
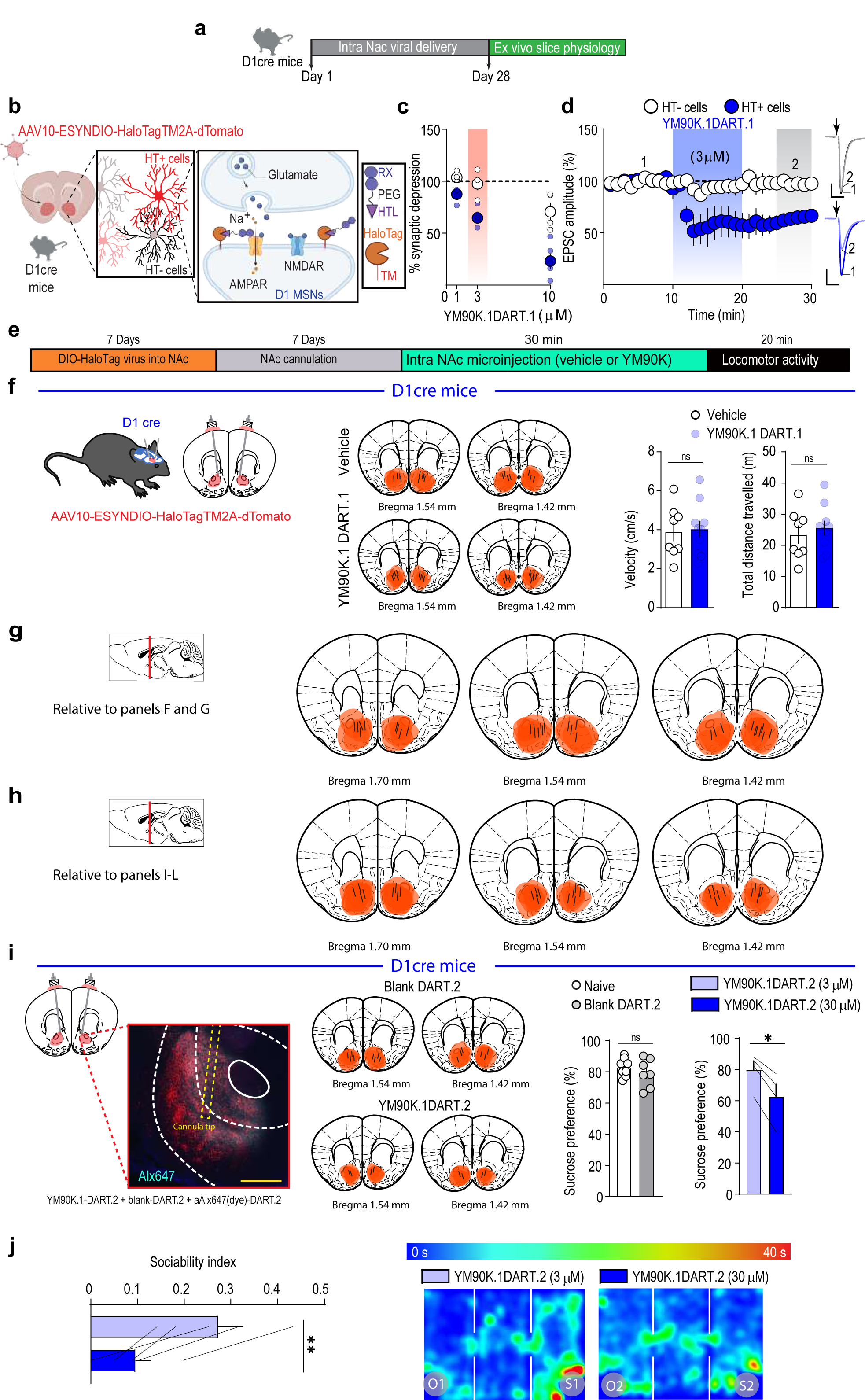
Validation of DART technology. **a**, **e**, Experimental timelines. **b**, Schematic of DART method. **c**, Synaptic depression in halotag- and + cells by bath application of YM90K.1^-DART.1^ ligand at different concentrations (1μM: *n* = 4 cells, 3 mice; 3μM: *n* = 5 cells, 3 mice; 10μM: *n* = 5 cells, 3 mice). **d**, Left: AMPAR-mediated synaptic depression induced with bath application of YM90K.1^-DART.1^ (3μM) in halotag- and + cells (halotag-: *n* = 5 cells, 3 mice; halotag+: *n* = 4 cells, 3 mice). Right, representative traces of YM90K.1^-DART.1^ bath application experiments. Scale bars, 20 ms, 200 pA. **f**, Left, schematic of the experiment. Center, mapping showing the expression of AAV10-ESYNDIO-HaloTagTM2A-dTomato and drawings illustrating locations of cannula tips in mice that received vehicle or YM90K.1^-DART.1^. Right, velocity and total distance travelled in D1-cre mice that received vehicle (*n* = 8) or YM90K.1^-DART.1^ (*n* = 10) microinjections in the NAc. **g**, **h**, Mapping showing the expression of AAV10-ESYNDIO-HaloTagTM2A-dTomato and drawings illustrating locations of cannula tips (**g**: relative to panels F and G in Fig. 2; **h**: relative to panels I and L in Fig. 2). **i**, Left, representative brain coronal section showing AAVrh10-ESYNDIO-HaloTagTM2A-dTomato transduction, cannula implantation track and YM90K.1^DART.2^ + Alexa488.1^DART.2^ in the NAc from a D1-cre mouse. Center, mapping showing the expression of AAVrh10-ESYNDIO-HaloTagTM2A-dTomato and drawings illustrating locations of cannula tips in mice that received Blank ^DART.2^ and YM90K.1^DART.2^. Right, sucrose preference in stress-naïve mice (*n* = 8) and stress-naïve mice that received Blank ^DART.2^ (*n* = 7) or YM90K.1^DART.2^ at 3μM and 30μM (*n* = 4) microinjections in the NAc. **J**, Left, sociability index in stress-naïve mice that received YM90K.1^DART.2^ at 3μM or 30μM microinjections in the NAc (*n* = 5). Right, representative occupancy plots from stress-naïve mice that received YM90K.1^DART.2^ at 3μM or 30μM microinjections in the NAc. Data are mean ± s.e.m. ns, not significant. \**P* < 0.05, \*\**P* < 0.01.

**Extended Data Fig. 5.**
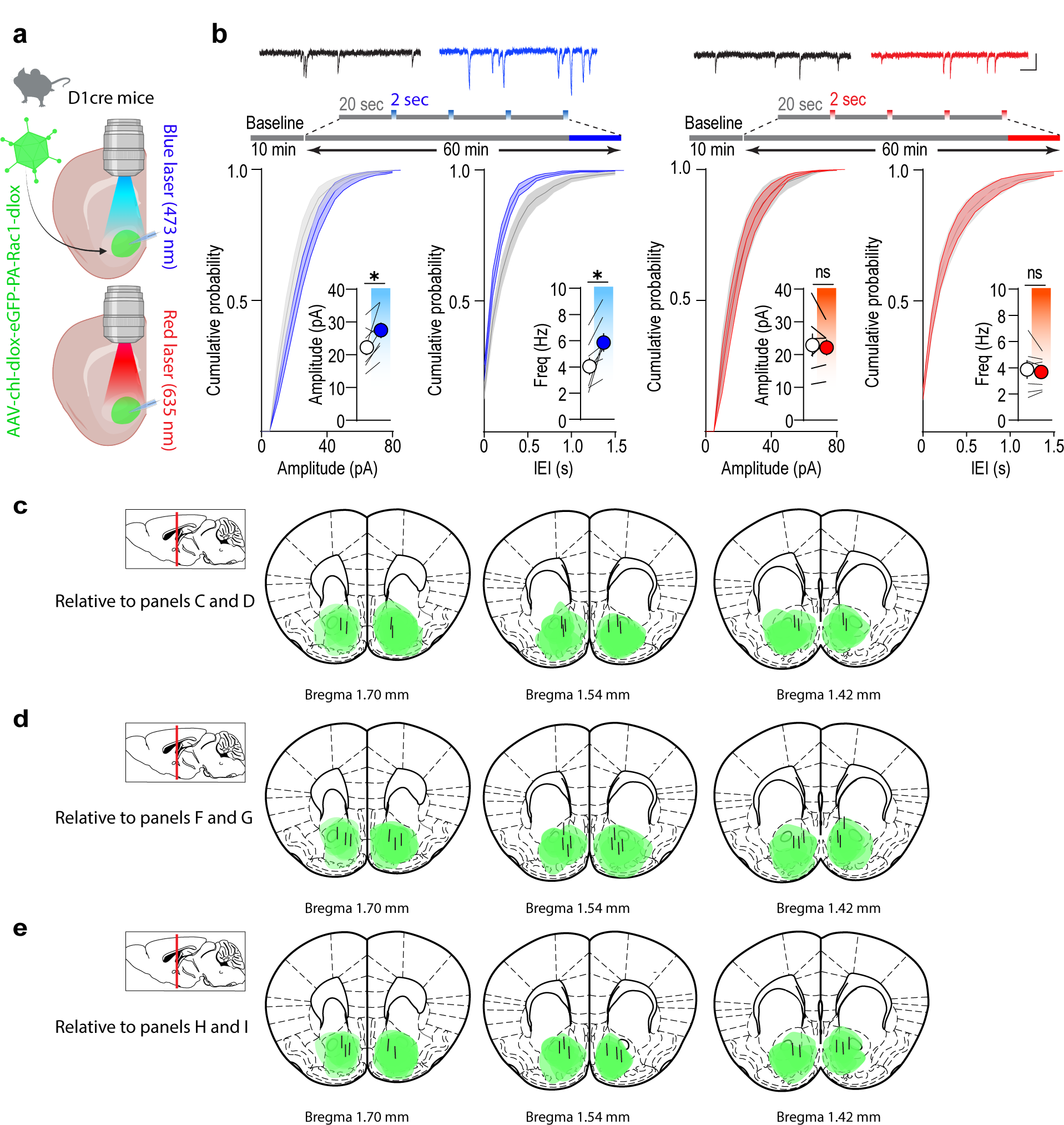
Validation of Pa-Rac1 methodology. **a,** Schematic depicting AAV-chl-dlox-eGFP-PA-Rac1-dlox injection and wavelengths used to stimulate the NAc MSNs of D1-cre mice in acute slice preparations. **b**, Top, example sEPSC traces and experimental protocol used for the *ex vivo* experiments. Scale bar, 100 ms, 20 pA. Bottom, cumulative probability plots of the amplitudes and frequencies of sEPSCs recorded from D1 cells in brain slices from stress-naïve mice before and after application of the blue (left) and red (right) lasers (for blue laser: *n* = 8 cells; 5 mice; for red laser: *n* = 7 cells; 5 mice). Insets: histograms of the means obtained from sEPSC amplitude and frequency. **c**, **d**, **e**, Mapping showing the expression of AAV-chl-dlox-eGFP-PA-Rac1-dlox and drawings illustrating locations of fiber tips (**c**: relative to panels C and D in Fig. 3; **d**: relative to panels F and G in Fig. 3; **e**: relative to panels H and I in Fig. 3). Data are mean ± s.e.m. ns, not significant. \**P* < 0.05.

**Extended Data Fig. 6.**
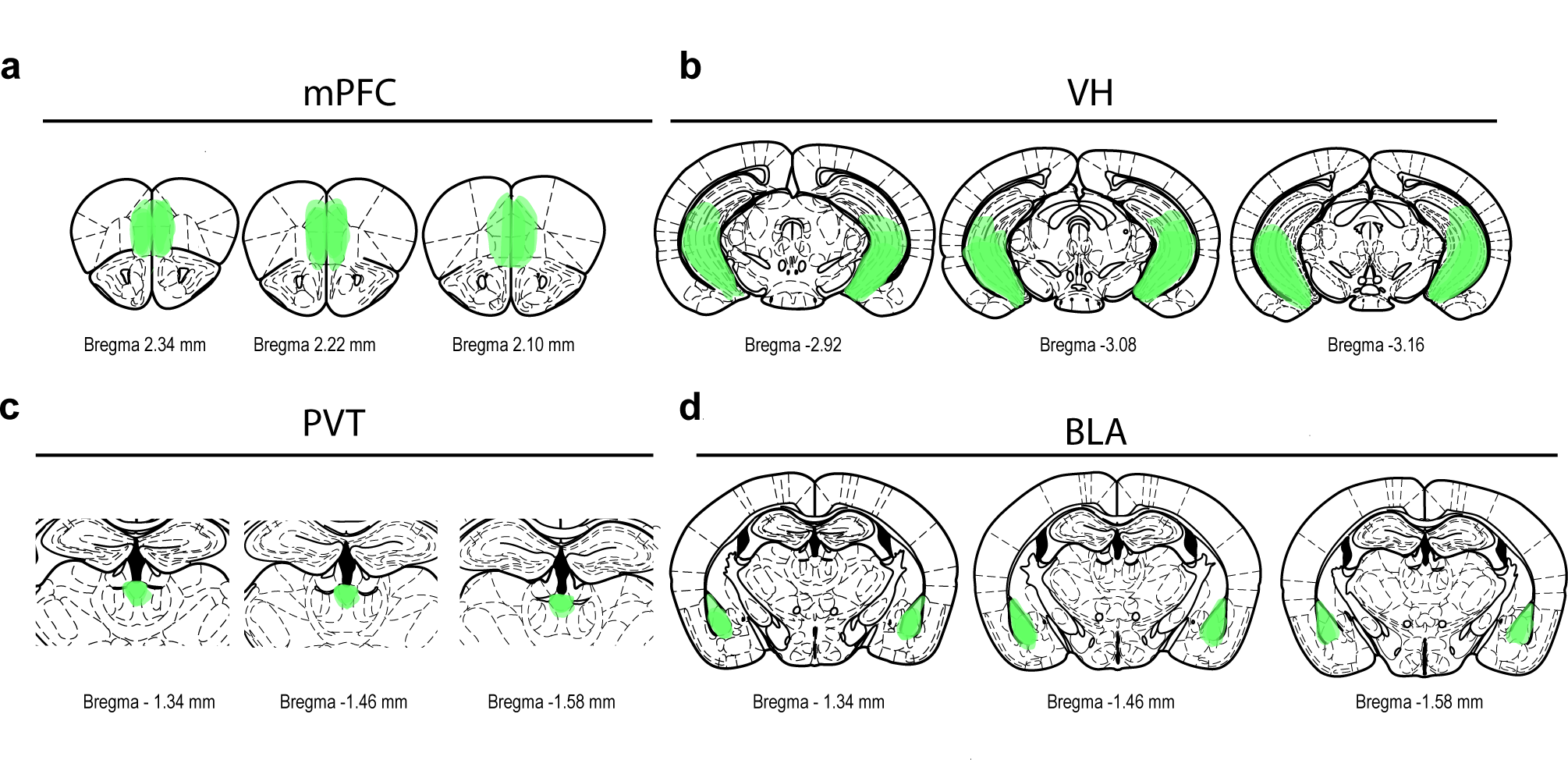
Pattern of expression of ChR2 in monosynaptic inputs to the NAc. Mapping showing the expression of AAV-camKII-ChR2-eYfp in mPFC (**a**), VH (**b**), PVT (**c**) or BLA (**d**).

**Extended Data Fig. 7.**
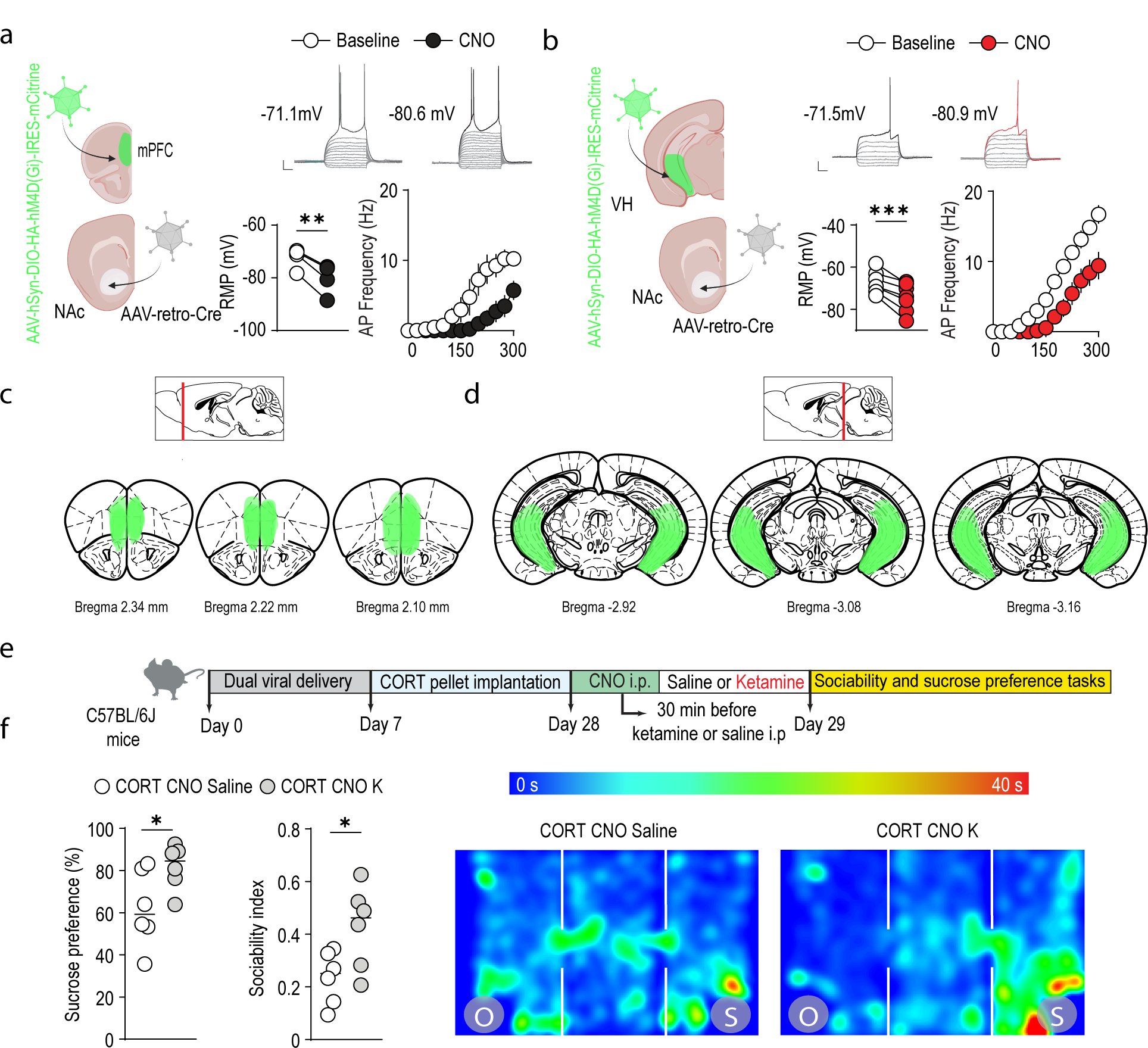
Validation of chemogenetic approach. **a, b,** Left, Schematic of the experiment. Right, resting membrane potential (RMP) and action potential (AP) frequency evoked by somatic depolarizing current steps before and after bath application of CNO (10 uM) from mPFC-(**a,** *n* = 4 cells; 2 mice) or VH-(**b,** *n* = 7 cells; 3 mice) NAc projecting pyramidal neurons. Scale bar, 100 ms, 10 mV. **c**, **d**, Mapping showing the expression of AAV-hSyn-DIO-HA-hM4D(Gi)-IRES-mcitrine in mPFC (**c**) or VH (**d**). **e**, Experimental timeline. **f**, Sucrose preference (left) and sociability index (center) in stressed mice that received CNO injections before saline (*n* = 6) or ketamine (*n* = 6) treatment. Right, representative occupancy plots from stressed mice that received CNO injections before saline or ketamine treatment. Data are mean ± s.e.m. \**P* < 0.05, ** *P* < 0.01.

**Extended Data Fig. 8.**
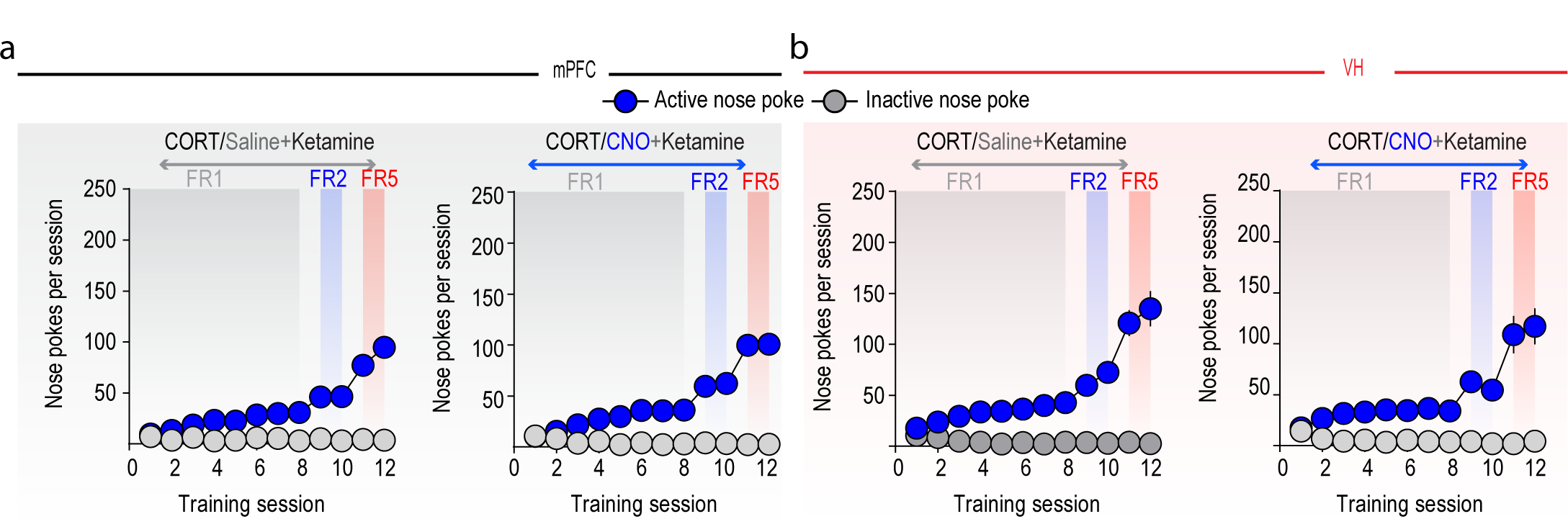
Sucrose self-administration acquisition in mice for the chemogenetic experiments. **a, b,** Performance during acquisition from stressed mice that received saline or CNO injections before ketamine treatment in the mPFC (**a**, saline: *n* = 12, CNO: n = 15) or VH (**b,** saline: *n* = 11, CNO: n = 10) groups.

## Notes

### Competing Interest Statement

The authors have declared no competing interest.

